# Species and intra-specific competition affect growth in greenhouse more than mycorrhizal colonization on *Quercus rubra* and *Acer rubrum* seedlings

**DOI:** 10.1101/2020.08.29.273300

**Authors:** Michael Maier

## Abstract

**Aims:** Distributions of mycorrhizal guilds correlate with differences in soil nutrient availability and tend to predominate ecosystems where plant growth and development are limited by either soil phosphorus (AM, arbuscular mycorrhiza) or nitrogen (EM, ectomycorrhiza) (Read, 1991). In an effort to characterize functional trait differences, we designed a common garden greenhouse experiment to measure growth and colonization responses between two tree seedling species (*Acer rubrum* and *Quercus rubra*) that associate with the two major mycorrhizal guilds.

**Methods:** In a fully factorial, 12-week greenhouse common garden experiment, seedlings were treated to three levels of nitrogen and phosphorus availability, and grown under inter- and intra-species competition. Change in heights and stem diameters were tracked and mycorrhizal colonization was quantified by percent of total and morphology.

**Results:** Relative growth rates were higher for *Acer rubrum* seedlings across treatments and intra-species competition had a strong negative effect on height and stem diameter, especially for *Quercus rubra*. Both species were highly colonized (>50%) by typically endomycorrhizal forms (arbuscules and vesicles) but varied in the distribution of morphological forms present in cells.

**Conclusions:** This study highlights the plasticity of tree seedling symbiosis during early developmental stages and challenges the strict static categorization of plant species associations with particular mycorrhizal guilds.

## Introduction

The mycorrhizal symbiosis is ubiquitous in vascular plants with about 85% forming some variety of fungal mutualism (M. C. Brundrett & Tedersoo, 2018). Most trees form either vesicular arbuscular mycorrhizae (AM) or ectomycorrhizal (EM) types of symbiosis, which are thought to be functionally linked to soil phosphorus (P) versus soil nitrogen (N) limitation respectively (Read, 1991). Differences in soil types and corresponding symbiotic fungi are characterized along a spectrum of environmental conditions where AM-associating species predominate in alkaline soils with relatively fast decomposition rates and low C:N ratios while EM plant hosts flourish where soil is more acidic, decomposition rates are slower, and C:N is low (Phillips, Brzostek, & Midgley, 2013; Read, 1991; Read & Perez-Moreno, 2003). This characterization can be thought of in terms of functional traits, where the more evolutionarily ancient AM types co-evolved with the development of vascular plant roots (Feijen, Vos, Nuytinck, & Merckx, 2018; Hoysted et al., 2018; Selosse & Tacon, 1998) and confer an advantage for plant development when accessing any nutrients, but especially limited phosphorus. The other two types of important mycorrhizae (EM and ericoid mycorrhizae) arose later through convergent evolution (Tedersoo & Smith, 2013), and recent evidence from evolution-based trait modelling (Lu & Hedin, 2019) highlights saprotrophic ancestors that were able to metabolize less accessible organic nitrogen sources through the production of hydrolytic and oxidative enzymes. Experimental results also underline this EM trait of organic N enzymatic processing (Bödeker et al., 2014) especially in those ectomycorrhiza descended from white rot as opposed to brown rot saprotrophic fungi (Stuart & Plett, 2020). Trait-based experimental design and analysis is increasingly recognized as a valuable method for characterizing guild differences (Lustenhouwer, Maynard, Bradford, Lindner, & Oberle, 2020; Mcgill, Enquist, Weiher, & Westoby, 2006; Zanne et al., 2020). Considering current projections of atmospheric CO_2_ increase, nitrogen deposition, phosphorus limitations, and the degradation and loss of forests worldwide (Crowther et al., 2016; Fitter, Heinemeyer, & Staddon, 2000; IPCC, 2014), clarifying the ecological function of mycorrhizal symbiosis is timely and relevant.

Studies comparing functional trait differences between EM and AM imply that we would expect to see a competitive advantage for EM trees under N limitation, and for AM trees under P limitation. This experiment was designed to test this using two tree species seedlings common to the Northeast US, across N and P gradients. AM and EM species often co-occur in New England forests, with some species sorting along moisture and geographic/soil gradients. Fossil records indicate that these two dominant tree genera in modern Northeastern US hardwood forests (*Acer* and *Quercus*, (Sapindaceae and Fagaceae)), evolved from different lineages and associate with different types of co-evolved mycorrhizae, endomycorrhizae or arbuscular mycorrhizae (AM) and ectomycorrhizae (EM) respectively (M. C. Brundrett & Tedersoo, 2018; Collins, Gostel, & Weeks, 2016; Robert, Mencuccini, & Martínez-Vilalta, 2017).

Seedlings are the critical life history stage for stand initiation and competition or cooperation within this stage is vital for the exclusion of other species and/or becoming dominant in a given location. Leveraging the benefits of mycorrhizal symbiosis early in developmental stages could represent a critical advantage for the long-term survival and dominance of seedlings within forests. We hypothesized that based on their mycorrhizal associations these two species should vary in their response to a nutrient gradient when grown in competition. By artificially creating N and P limitation in the soil, we could then provide a better understanding of the competitive advantages of AM and EM inoculated seedlings in terms of biomass in the seedling life history stage. Biomass accumulation in terms of relative growth rates and percent fine root colonization were used as metrics for characterizing how functional trait differences in mycorrhizal guilds affect tree seedling competition. Based on functional trait differences, we tested the following hypotheses: (i) Their dominance and high distribution in phosphorus-poor soils suggest that arbuscular mycorrhiza (AM) associated seedlings (*Acer rubrum L.*) will compete favorably where soil phosphorus is limited when in direct competition with (ii) ectomycorrhizal species (EM, *Quercus rubra L.*), who should compete more favorably where soil nitrogen is limited.

## Materials and Methods

Seedling competition was included as a factor (Figure 1) to test our hypothesis that different mycorrhizal guilds provide functional trait advantages according to variations in nutrient availability. Treatment and species mix were replicated five times in a fully factorial balanced common garden design. Ninety *Acer rubrum* seedlings and 90 *Quercus rubra* seedlings were grown in monoculture and combination and treated with one of three nutrient solutions twice a week for 3 months. While our focus was the contrast provided by interspecific competition treatments, we were also interested in responses when the same nutrient variation was applied to monocultures.

**Figure 1.**
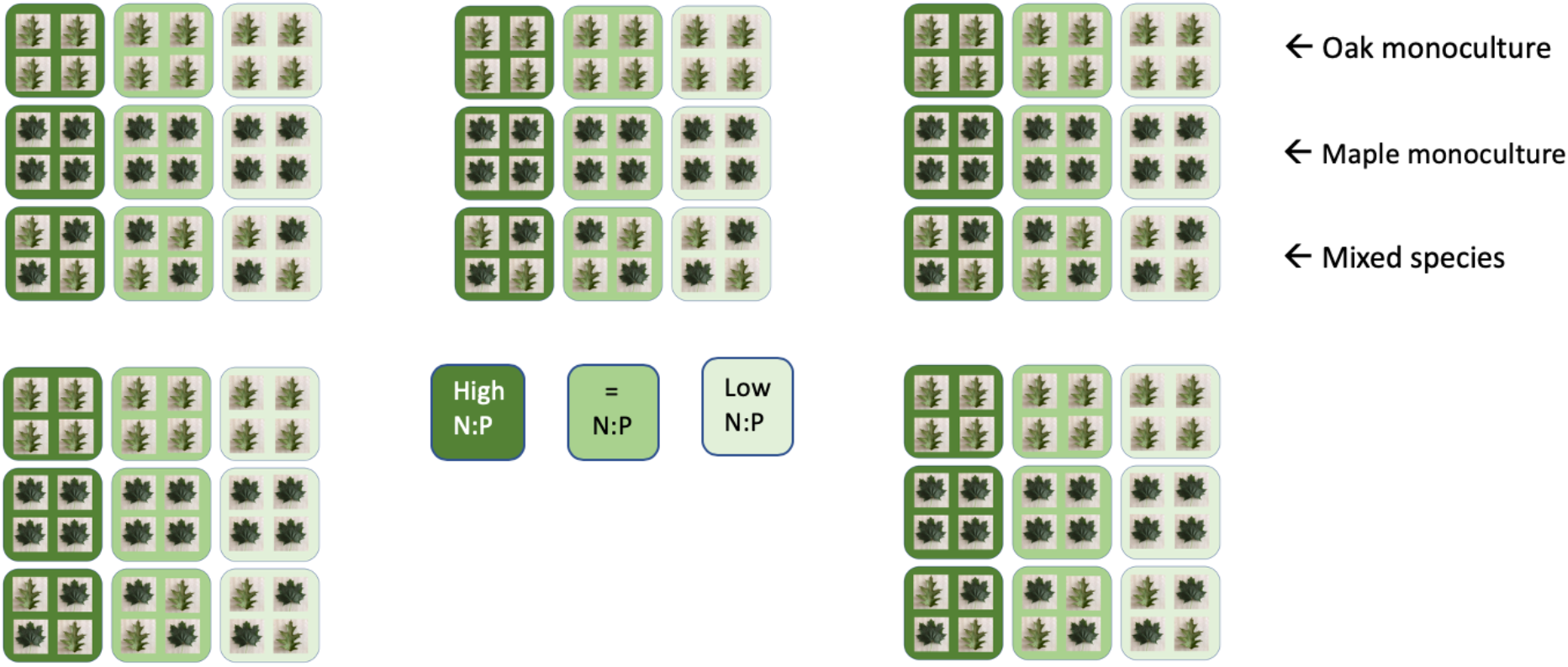
Treatment and species mix replicated five times in a fully factorial balanced common garden design. 90 red maple seedlings and 90 red oak seedlings were grown in monoculture and combination (Intra and Inter species competition) and treated with one of three nutrient solutions twice a week for 3 months.

Thirty-centimeter average size bare-root *Acer rubrum* and *Quercus rubra* seedlings (Prides Corner Nursery, Lebanon CT) were grown in potting soil for three months before treatment. At the start of the experiment, roots were replanted into 10L pots with sterilized soil mix (40% sterilized sand/ 60% Griffins Supplies BK55ProMix-a combination of processed southern pine bark, sphagnum peat moss, and vermiculite) and grown in a greenhouse. Supplemental lighting and heat were provided to maintain average summer temp (21.5 C) and photoperiod (15hr light/9hr dark). In a fully factorial (2 × 3 × 2) greenhouse experiment the two species of seedlings were provided with associated mycorrhiza field inoculant, and were treated at 3 levels of fertilization (Low N:P, Balanced N:P, High N:P) and two levels of competition: intraspecific and interspecific.

Considering the importance of co-adapted mycorrhizal species in promoting growth (Johnson, Wilson, Bowker, Wilson, & Miller, 2010), seedlings were inoculated with fine roots collected from established *Acer rubrum* and *Quercus rubra* locations within 48 hours of collection (Orchard, Standish, Nicol, Dickie, & Ryan, 2017). Unidentified field mixtures of fine root material were isolated from 1kg soil samples from Yale Myers Forest in Union, CT (41.965729, −72.125162) cleaned by repeated rinsing, mixed with 2.7l DI water, and homogenized in a blender to produce 30ml of solution per seedling. Each of the single donor trees growing in the fine root source had similar dbh (between 50-60 cm), crown size, and topographic environment and were located within 250 m of each other. At the outset of treatments, we administered 30ml of species-specific inoculant to each seedling.

Balanced nutrient solution with a full complement of macro and micronutrients served as the control baseline for growth response and followed the basal salt solution as outlined by (Murashige & Skoog, 1962) containing inorganic salts NH_4_NO_3_ (400mg/l), KCl (65mg/l), KNO_3_ (80mg/l), KH_2_PO_4_ (12.5mg/l), Ca(NO_3_)_4_ ·4H_2_O (144mg/l), MgSO_4_ ·7H_2_O (72mg/l), NaFe-EDTA (25mg/l), H_3_BO_3_ (1.6mg/l), MnSO_4_ · 4H_2_O (6.5mg/l), ZnSO_4_ ·7H_2_O (2.7mg/l), and KI (0.75mg/l). High N:P solution was the same basal mix but without potassium phosphate monbasic and Low N:P was the same basal mix but without ammonium nitrate and potassium nitrate. We followed a similar protocol to Holste & Kobe (2017), and applied 10ml of the nutrient solutions twice per week to each pot.

Each pot was randomized within a grid/matrix on the greenhouse bench under supplemental lighting and automated irrigation (at an average rate of 2L/hour for 10 min/day). These conditions provided for well-watered seedlings to avoid confounding effects from drought stress and allowed for observation of growth resulting only from variations in nutrients and the influence of mycorrhiza. Soil moisture was monitored to measure any variation in water utilization with a Campbell (HS2) handheld soil water sensor. Irrigation was provided to keep all soil at or near 50% of field capacity. During week 8, aphid infestation of more than 50% of all seedlings was successfully treated with two applications of Suffoil-X (a spray oil emulsion insecticide that is 80% mineral oil).

Seedling heights were measured weekly, stem diameter was recorded with calipers pre- and post-treatment, and symbiotic development (% root colonization) was measured after 12 weeks. Field soil N and P levels (Table 1) were tested and compared to the experimental growth medium. The experimental soil medium mix of sterile sand and potting soil had comparable pH (5.5) and available Nitrate (3ppm), but had less available Ammonium (12ppm) and more available P (100 ppm). *Quercus* root samples were initially examined at 40x under a dissecting microscope following the ectomycorrhizal quantification methods of (M. Brundrett et al., 1996). Further post treatment analysis of percent mycorrhizal colonization for both species was ascertained through selective root staining and the compound microscopy intersection methods at higher magnification (200x and 500x) outlined for endomycorrhizal samples in (M. Brundrett et al., 1996; Dobson, Richardson, & Blossey, 2020; West Virginia University, 2019).

**Table 1.** Soil nutrient availabilities. Potting soil indicates the medium in which the seedlings grew up to 3 months. Available macronutrient amounts are listed in ppm. Experiment represents the mix of sterile potting soil and sterilized sand. *Acer* and *Quercus* are samples from the donor trees that supplied material for inoculants. Experimental mix had lower levels of available N, but higher levels of available P than inoculant parent soils.

| Soil | Texture | Organic Matter | pH | Nitrate N (ppm) | Ammonium N (ppm) | Phosphorus (ppm) | Calcium (ppm) | Magnesium (ppm) |
| --- | --- | --- | --- | --- | --- | --- | --- | --- |
| Potting | Organic | Very High | 6.0 | 25 | 150 | 75 | 900 | 18 |
| Experiment | Mix | Very High | 5.5 | 3 | 12 | 100 | 900 | 80 |
| Acer | Sandy Loam | Medium | 5.4 | 3 | 105 | 19 | 700 | 18 |
| Quercus | Sandy Loam | Medium | 5.1 | 3 | 56 | 19 | 700 | 25 |

### Root staining and histology

At the conclusion of treatments, roots were soaked and rinsed in water until all sand and soil was removed, and 0.1g samples of fine root tissue were packed loosely in biopsy cassettes with 0.9mm holes. Four 0.1g samples were taken from each seedling. To sufficiently clear seedling roots, cassettes were autoclaved two at a time in 100ml of 20% KOH solution for 30 minutes at 121C on the liquids cycle. After rinsing thoroughly with DI water, cassettes were soaked at room temperature in 6% H_2_O_2_ for at least 2 hours, and up to 4 hours depending on how translucent the tissue had become. After a 5-minute immersion in 2% HCl, samples were soaked overnight in staining solution of 6ml toluidine blue:450ml glycerol:450ml DI water. After at least 24 hours at room temperature, samples were rinsed and stored in 1:1 DI water and glycerol for a further 24 hours before mounting.

Twenty 1cm sections were taken from each sample cassette and mounted on standard slides with #1.5 (0.17mm) coverslips. Pressure was applied by hand with an extra slide wrapped in a Kimwipe for about 30 seconds to flatten the root tissue. 100 intersections per slide (5 per section) were recorded using an Olympus BX60 compound microscope (at 200x and 500x) and categorized as 1-root only, 2-hyphae only, 3-arbuscules and hyphae, and 4-vesicles and hyphae.

### Statistics

Growth in terms of change in both stem diameter and seedling height over time were calculated as Relative Growth Rate where S2 is final growth measure, S1 is initial growth measure and t2 and t1 are final time and initial time:

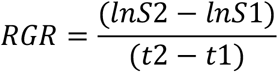

Data sets were checked for linearity, homogeneity of variance, and normal distribution of residuals. Differences among group treatment means were modelled with analysis of variance and post-hoc Tukey’s HSD was performed in R with the *agricolae* package, (Mendiburu, 2020). Interactions between treatments, competition level, growth metrics, and colonization levels were further analyzed with mixed effects linear models using the *lme4* package (Bates, Mächler, Bolker, & Walker, 2015) and visualized with *multcompView* (Graves, Piepho, Selzer, & Dorai-Raj, 2019) and *sjPlot* (Lüdecke, 2020). Post-hoc Tukey tests on mixed linear effects models were made with *emmeans* (Lenth, 2020). For mixed effects models where stem diameter and height were dependent variables, species, treatment, and competition type were modelled as additive predictor variables with unit (a designation of one of the 5 repetitions of the 9 different treatment combinations, Figure 1) as a random effect. Where colonization percentage was the dependent variable, species, treatment, their interaction, and competition type were modelled as additive predictor variables with unit as a random effect. To analyze effect size for both ANOVA and mixed linear effects models, omega squared (ω^2^) and eta squared (η^2^) were calculated in addition to r^2^. Eta squared is an indication of the amount of variance associated with each effect (Lakens, 2013), while omega squared is an additional indication of the strength of the relationship between independent variables and responses that applies especially to relatively small sample sizes (Albers & Lakens, 2018; Field, 2013). Suggested effect value ranges for omega squared are very small (0-0.01), small (0.01-0.06), medium (0.06-0.14), and large (>0.14). Each of these effect size statistics helps distinguish between when a response is statistically significant and also has substantial impact versus statistical significance but insubstantial impact. All statistical analysis and data visualization was done in R Version 3.6.2 utilizing *tidyverse* (Wickham et al., 2019) and *ggplot2* (Wickham, 2009).

## Results

### Growth Response

In terms of stem diameter change over time, analysis of variance showed that species had the largest and most significant effect (*F*_*1,168*_ = 45.732, *p* < 0.001, *r*^*2*^= 0.27). Effect size in terms of omega squared (ω^2^ = 0.20) and eta squared (η^2^ = 0.21) were substantial. *Quercus rubra* seedlings showed significantly less change over time than *Acer rubrum* seedlings regardless of treatment or competition level (Figure 2). Change in seedling heights were mainly influenced by species (*F*_*1,168*_ = 20.354, *p* < 0.001, *r*^*2*^= 0.27) and competition (*F*_*1,168*_ = 34.240, *p* < 0.001, *r*^*2*^= 0.27); *Acer rubrum* species and interspecies competition demonstrated significantly more growth than *Quercus rubra* and intraspecies competition. Effect size in terms of Omega squared (ω^2^ = 0.10 for Species and 0.16 for Competition) was medium to large according to the heuristics of (Field, 2013).

**Figure 2.**
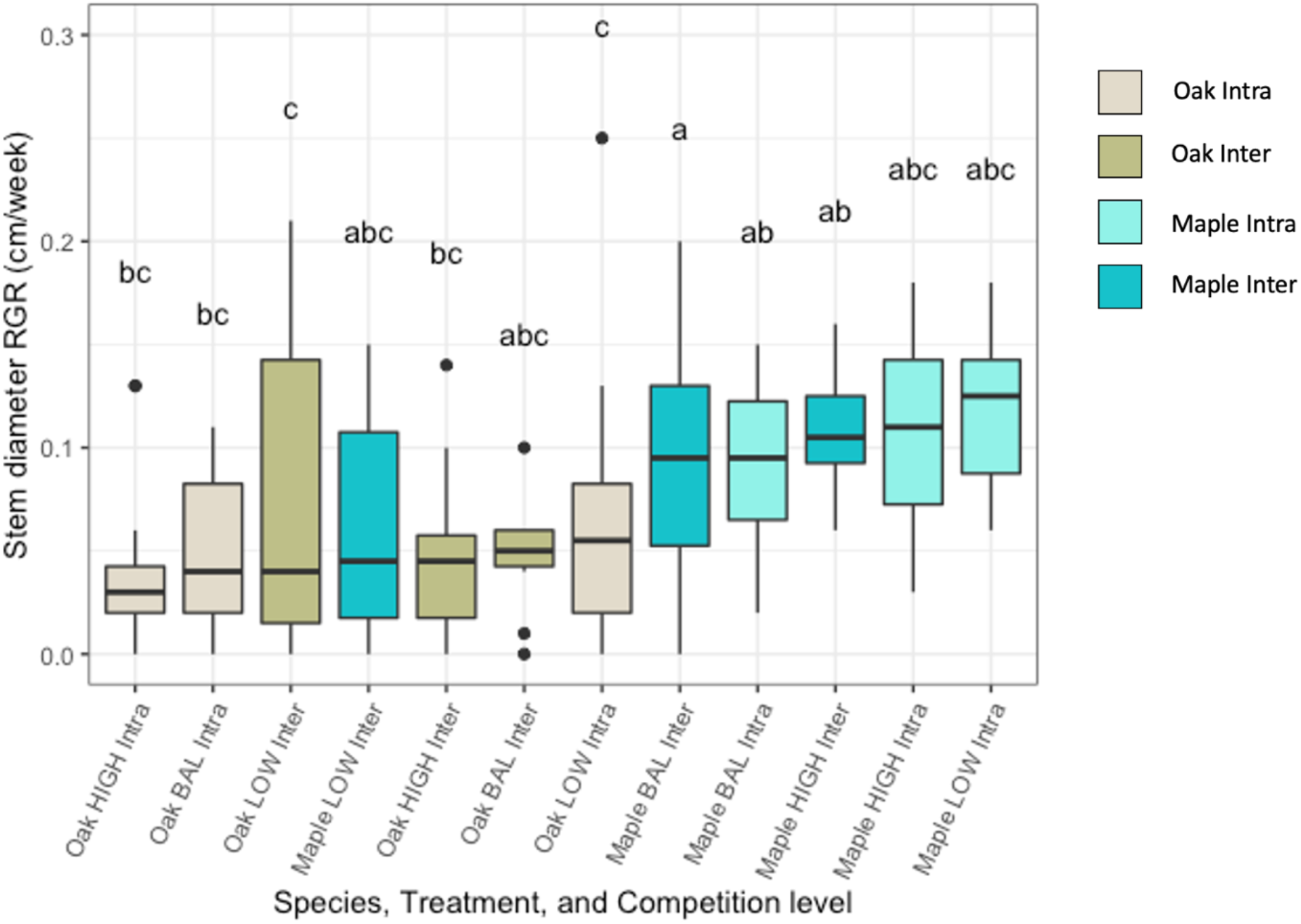
Stem diameter change over time by species and nutrient treatment when *Acer rubrum* and *Quercus rubra* seedlings are grown in both monoculture and in mixed competition for 3 months. Lower box boundary indicates the 25th percentile, the median value is shown as black line, and the upper boundary contains the 75th percentile. Error bars are whiskers above and below showing the 90th and 10th percentiles. Points outside the 90th and 10th percentiles (outliers) are also represented. Plots with the *same letters* above each box do not differ significantly among treatment groups.

A linear mixed effects model for relative growth rate changes in seeding heights (left side of Table 2) point out significant differences in means (*p* < 0.001, *r*^*2*^ = 0.33) between species as well as between intra- and interspecies competition. For species, this effect size is not large (ω^2^ = 0.11, (η^2^ = 0.12) but for competition the effect size is substantial (ω^2^ = 0.35, (η^2^ = 0.37). A linear mixed effects model for stem diameter growth rate (right side of Table 2) shows species as significant (*p* < 0.001, *r*^*2*^= 0.27, (ω^2^ = 0.19, (η^2^ = 0.20). Tukey post hoc HSD supports significant species differences.

**Table 2.**
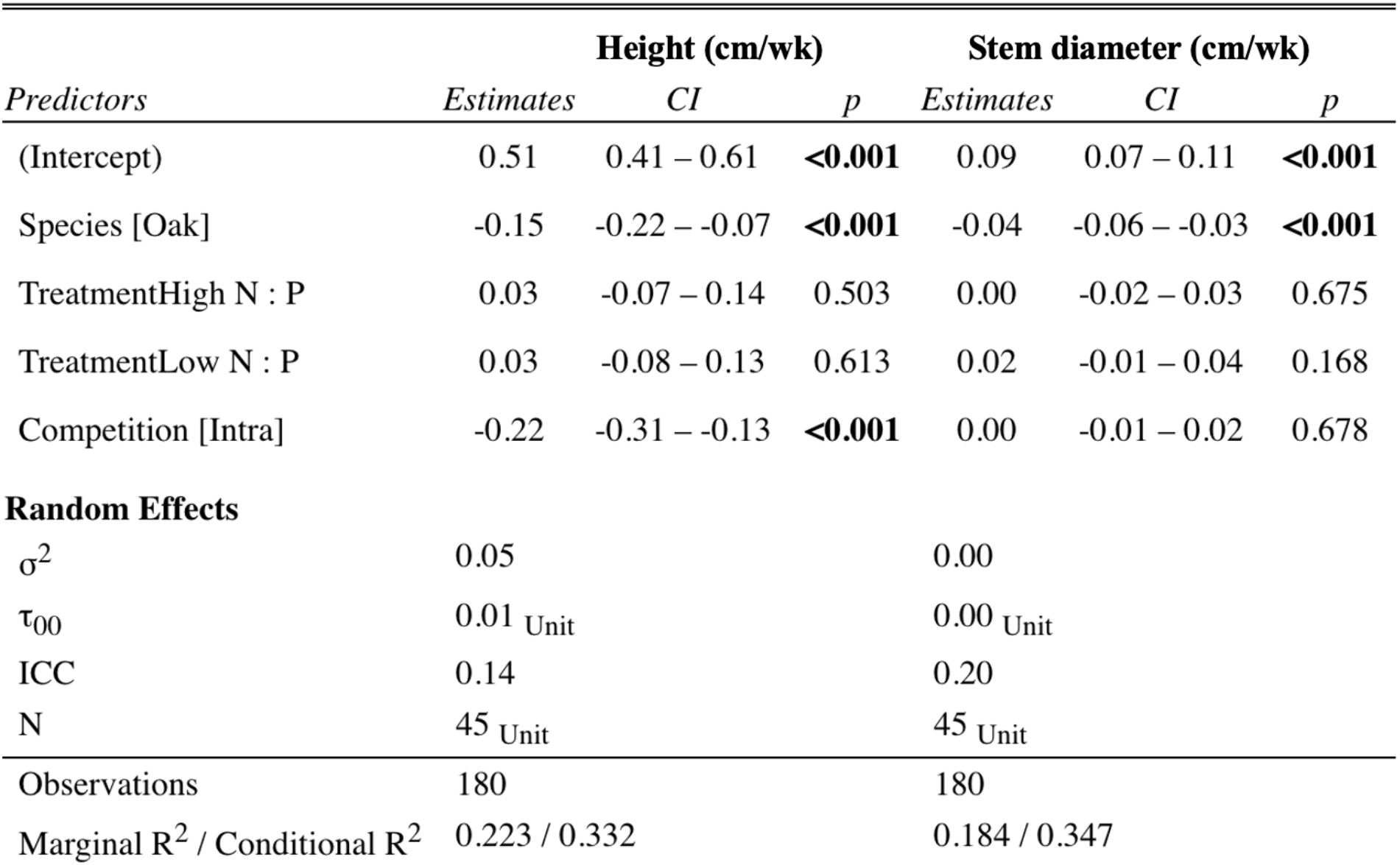
Linear mixed effects model results for Relative Growth Rate for changes in seedling heights and stem diameter. The lmer function automatically calculates t-tests using Satterthwaite approximations to degrees of freedom. 95% confidence intervals were calculated for means. Marginal R^2^ a measure of the proportion of variability explained by the fixed effects. Conditional R^2^ is the proportion of total variance explained through both fixed *and* random effects.

| *Predictors* | **Height (cm/wk)** |  |  | **Stem diameter (cm/wk)** |  |  |
| --- | --- | --- | --- | --- | --- | --- |
|  | *Estimates* | *CI* | *p* | *Estimates* | *CI* | *p* |
| (Intercept) | 0.51 | 0.41 – 0.61 | **<0.001** | 0.09 | 0.07 – 0.11 | **<0.001** |
| Species [Oak] | -0.15 | -0.22 – -0.07 | **<0.001** | -0.04 | -0.06 – -0.03 | **<0.001** |
| TreatmentHigh N : P | 0.03 | -0.07 – 0.14 | 0.503 | 0.00 | -0.02 – 0.03 | 0.675 |
| TreatmentLow N : P | 0.03 | -0.08 – 0.13 | 0.613 | 0.02 | -0.01 – 0.04 | 0.168 |
| Competition [Intra] | -0.22 | -0.31 – -0.13 | **<0.001** | 0.00 | -0.01 – 0.02 | 0.678 |
| **Random Effects** |  |  |  |  |  |  |
| $\sigma^2$ | 0.05 | | | 0.00 | | |
| $\tau_{00}$ | 0.01 Unit | | | 0.00 Unit | | |
| ICC | 0.14 |  |  | 0.20 |  |  |
| N | 45 Unit |  |  | 45 Unit |  |  |
| Observations | 180 |  |  | 180 |  |  |
| Marginal R<sup>2</sup> / Conditional R<sup>2</sup> | 0.223 / 0.332 |  |  | 0.184 / 0.347 |  |  |

### Mycorrhizal Response

Evidence for the expected ectomycorrhizal (EM) colonization of *Quercus* seedlings was absent in fine root tissue samples in 3-month-old seedlings after 12 weeks of treatment and development. Endomycorrhizal (AM) development however, was present in both species mostly in the form of extensive vesicles in hypodermis and cortex cells as outlined in (M. Brundrett et al., 1996) of *Quercus rubra* (Figure 4) and arbuscules and extensive hyphae in *Acer rubrum* seedlings (Figure 5). Colonization levels were > 50% across all species and treatments (Figure 6).

Colonization percentages were not significantly affected by treatment, species, or competition, however there was a significant interaction between nutrient treatment and species (*F*_*2,168*_ = 7.224, *p* <0.001, *r*^*2*^ = 0.16, ω^2^ = 0.06, η^2^ = 0.08). Interaction between treatment and competition (*F*_*2,168*_ = 3.280, *p* = 0.04, *r*^*2*^ = 0.16, ω^2^ = 0.06, η^2^ = 0.08) and interaction between species and competition (*F*_*1,168*_ = 0.558, *p* = 0.042, *r*^*2*^ = 0.16, ω^2^ = 0.06, η^2^ = 0.08) were also highlighted. Low N:P *Acer* vs. Balance *Acer* (*p* = 0.002) and Balance *Acer* vs. Balance *Quercus* (*p* = 0.03) were the two main combinations highlighted in post-hoc Tukey tests.

Fungal morphology (presence of arbuscules and hyphae, vesicles and hyphae, and hyphae only) varied substantially between species. Total colonization percentage (Table 3), was explained well by our model (*r*^*2*^ = 0.79) and indicates (*p* = 0.001) that *Quercus* has higher colonization, though effect size is small (ω^2^ = 0.06, η^2^ = 0.06). The type of mycorrhizal form varied distinctly between species (*p* < 0.001) in highly explanatory models (Table 3, *r*^*2*^ > 0.87). *Acer rubrum* had particularly high rates of hyphae with pronounced effect size (ω^2^ = 0.66, η^2^ = 0.66). *Quercus rubra* showed a distinct difference in incidences of arbuscules (*p* < 0.001, *r*^*2*^ = 0.91) though the size of the effect was not particularly large (ω^2^ = 0.37, η^2^ = 0.38). *Quercus* also showed a significantly higher number of vesicles (*p* < 0.001, *r*^*2*^ = 0.95) with large effect sizes (ω^2^ = 0.85, η^2^ = 0.85). Low N:P was highlighted in terms of vesicle formation though at a lower significance (*p* =0.04) and with a small effect size (ω^2^ = 0.10, η^2^ = 0.06). Pearson’s chi square tests of mycorrhizal type subset by species shows no associations between a particular treatment and count of forms (i.e., arbuscules, vesicles, hyphae).

**Table 3.** Results of linear mixed effects models of different mycorrhizal morphology in terms of total colonization percent, arbuscules, vesicles, and hyphae between species, treatments, and competition levels. The lmer function automatically calculates t-tests using Satterthwaite approximations to degrees of freedom. 95% confidence intervals were calculated for means. Marginal R^2^ a measure of the proportion of variability explained by the fixed effects. Conditional R^2^ is the proportion of total variance explained through both fixed *and* random effects.

| Predictors | % colonization |  |  | hyphae |  |  | hyphae and arbuscules |  |  | hyphae and vesicles |  |  |
| --- | --- | --- | --- | --- | --- | --- | --- | --- | --- | --- | --- | --- |
|  | Estimates | CI | p | Estimates | CI | p | Estimates | CI | p | Estimates | CI | p |
| (Intercept) | 59.30 | 50.22 – 68.38 | **<0.001** | 46.86 | 38.80 – 54.92 | **<0.001** | 8.29 | 2.91 – 13.67 | **0.003** | 4.04 | -2.57 – 10.65 | 0.231 |
| treatmentHigh N : P | -1.44 | -11.21 – 8.33 | 0.773 | -1.80 | -10.43 – 6.83 | 0.683 | -1.47 | -7.31 – 4.38 | 0.623 | 1.83 | -5.31 – 8.98 | 0.615 |
| treatmentLow N : P | 4.32 | -5.45 – 14.09 | 0.386 | -3.73 | -12.36 – 4.89 | 0.396 | 0.50 | -5.34 – 6.34 | 0.867 | 7.57 | 0.42 – 14.71 | **0.038** |
| species [Oak] | 5.75 | 2.37 – 9.12 | **0.001** | -32.50 | -35.98 – -29.02 | **<0.001** | 6.67 | 5.29 – 8.05 | **<0.001** | 31.79 | 29.68 – 33.90 | **<0.001** |
| competition [Intra] | 1.86 | -6.60 – 10.32 | 0.667 | 4.33 | -3.14 – 11.80 | 0.256 | -0.23 | -5.29 – 4.83 | 0.928 | -2.20 | -8.39 – 3.99 | 0.486 |
| **Random Effects** |  |  |  |  |  |  |  |  |  |  |  |  |
| $\sigma^2$ | 50.59 | | | 56.50 | | | 7.89 | | | 18.98 | | |
| $\tau_{00}$ | 173.73<sub>unit</sub> | | | 131.12<sub>unit</sub> | | | 64.67<sub>unit</sub> | | | 94.96<sub>unit</sub> | | |
| ICC | 0.77 |  |  | 0.70 |  |  | 0.89 |  |  | 0.83 |  |  |
| N | 45<sub>unit</sub> |  |  | 45<sub>unit</sub> |  |  | 45<sub>unit</sub> |  |  | 45<sub>unit</sub> |  |  |
| Observations | 180 |  |  | 180 |  |  | 180 |  |  | 180 |  |  |
| Marginal R<sup>2</sup> / Conditional R<sup>2</sup> | 0.063 / 0.789 |  |  | 0.592 / 0.877 |  |  | 0.141 / 0.907 |  |  | 0.700 / 0.950 |  |  |

## Discussion

Generally, seedling growth did not support the functional trait differences between AM- and EM-associating species we hypothesized and variations in available nutrients did not consistently or significantly spur growth in species. Low N treatments for *Quercus* and Low P treatments for *Acer* did not affect growth in terms of either seedling heights or stem diameters. Interestingly, both *Quercus rubra* and *Acer rubrum* seedlings were colonized by endomycorrhizal forms (AM). Differences in stem diameters (Figure 2) and seedling heights (Figure 3) varied mainly by species and by whether the plants were grown in monoculture or competed with the other study species. Our results showed that while variations in nutrient treatments did not affect growth during this period, seedling species and the type of competition those seedlings face makes more of a difference in terms of growth and development. Greater heights under interspecific competition stand out clearly (Figure 3). The mechanism behind this appears to have little to do with available nutrients or the type of species as seedlings grow more in terms of height much less when they are in monoculture and significantly more when they are in competition with another species. Physiologically, there are distinct differences between the species (*Acer* and *Quercus*) that explain variation in growth rates (Comas & Eissenstat, 2004; Lorimer, 1984; Medeiros, Tomeo, Hewins, & Rosenthal, 2016) but why does interspecific competition result in greater heights? Our results support the contention that intraspecific competition has a significant negative impact with a larger effect than interspecific competition (Adler et al., 2018).

**Figure 3.**
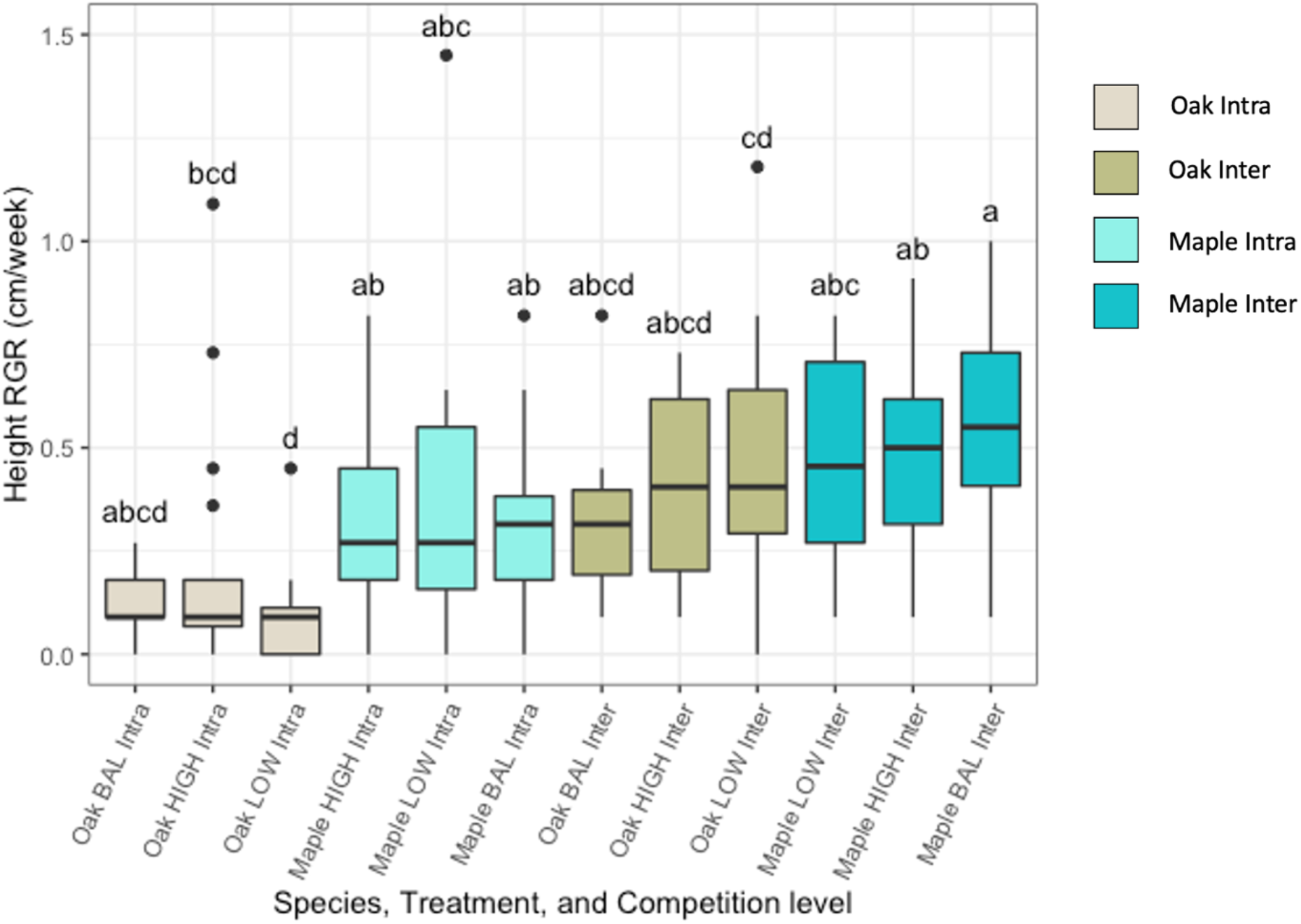
Seedling heights by species and nutrient treatment when *Acer rubrum* and *Quercus rubra* seedlings are grown in both monoculture and in mixed competition for 3 months. Lower box boundary indicates the 25th percentile, the median value is shown as black line, and the upper boundary contains the 75th percentile. Error bars are whiskers above and below showing the 90th and 10th percentiles. Points outside the 90th and 10th percentiles (outliers) are also represented. Plots with the *same letters* above each box do not differ significantly by ANOVA (*p* < 0.05) among treatment groups.

While the source soil used for inoculation did contain evidence of EM (i.e. fine roots clearly showed mantle formation), *Quercus* roots were uncolonized by ectomycorrhizas. Both species were highly colonized by typically AM forms under both competition levels and across treatments (Figures 4 and 5). Explanations for this could include pre-inoculation infection as nursery seedlings or the presence of competitive AM species in fine roots from *Quercus* stands used for the seedling inoculant.

**Figure 4.**
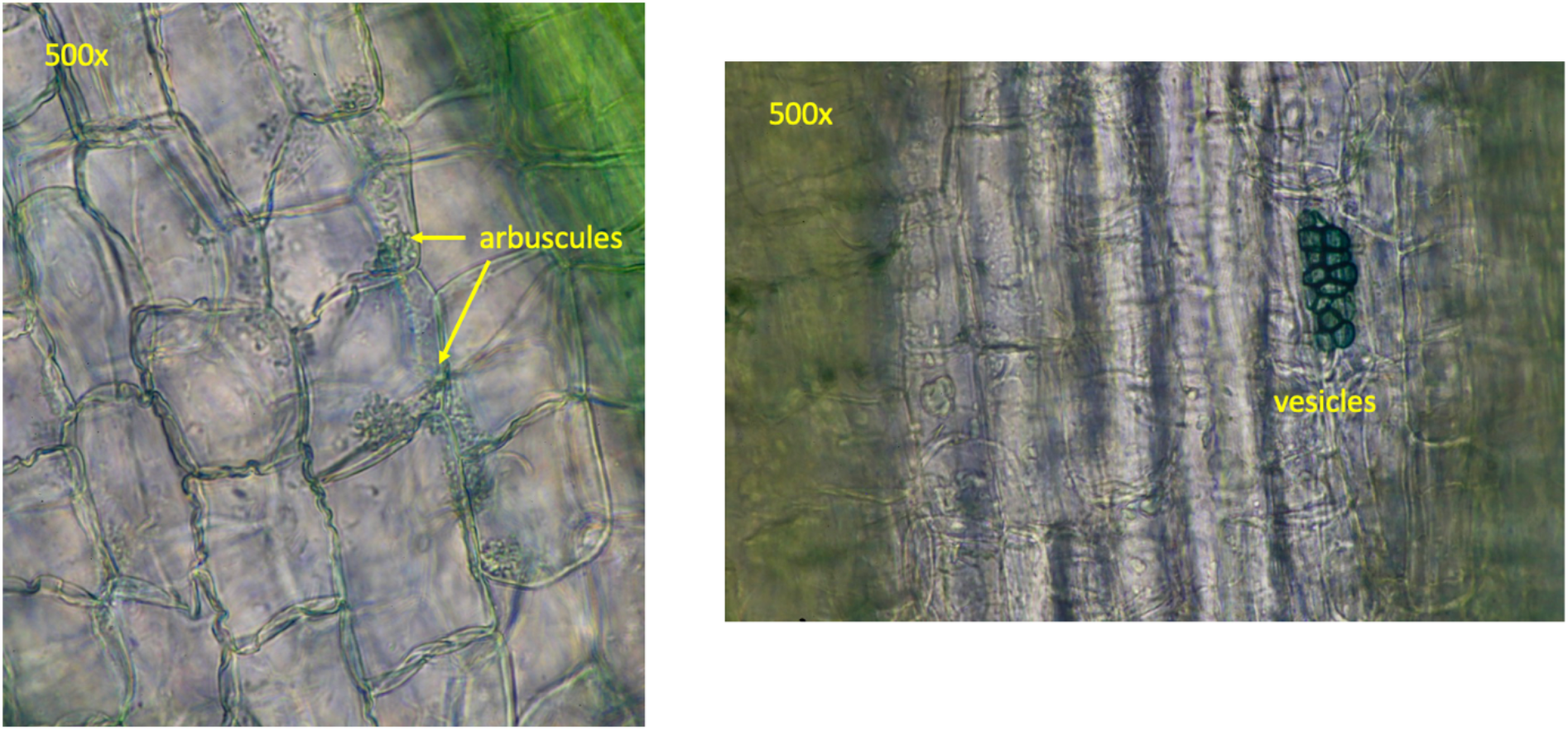
Compound microscope images (Olympus BX-60) of cleared and stained *Quercus rubra* root tissue showing the development of both arbuscules and vesicles typical of endomycorrhizal colonization (AM).

**Figure 5.**
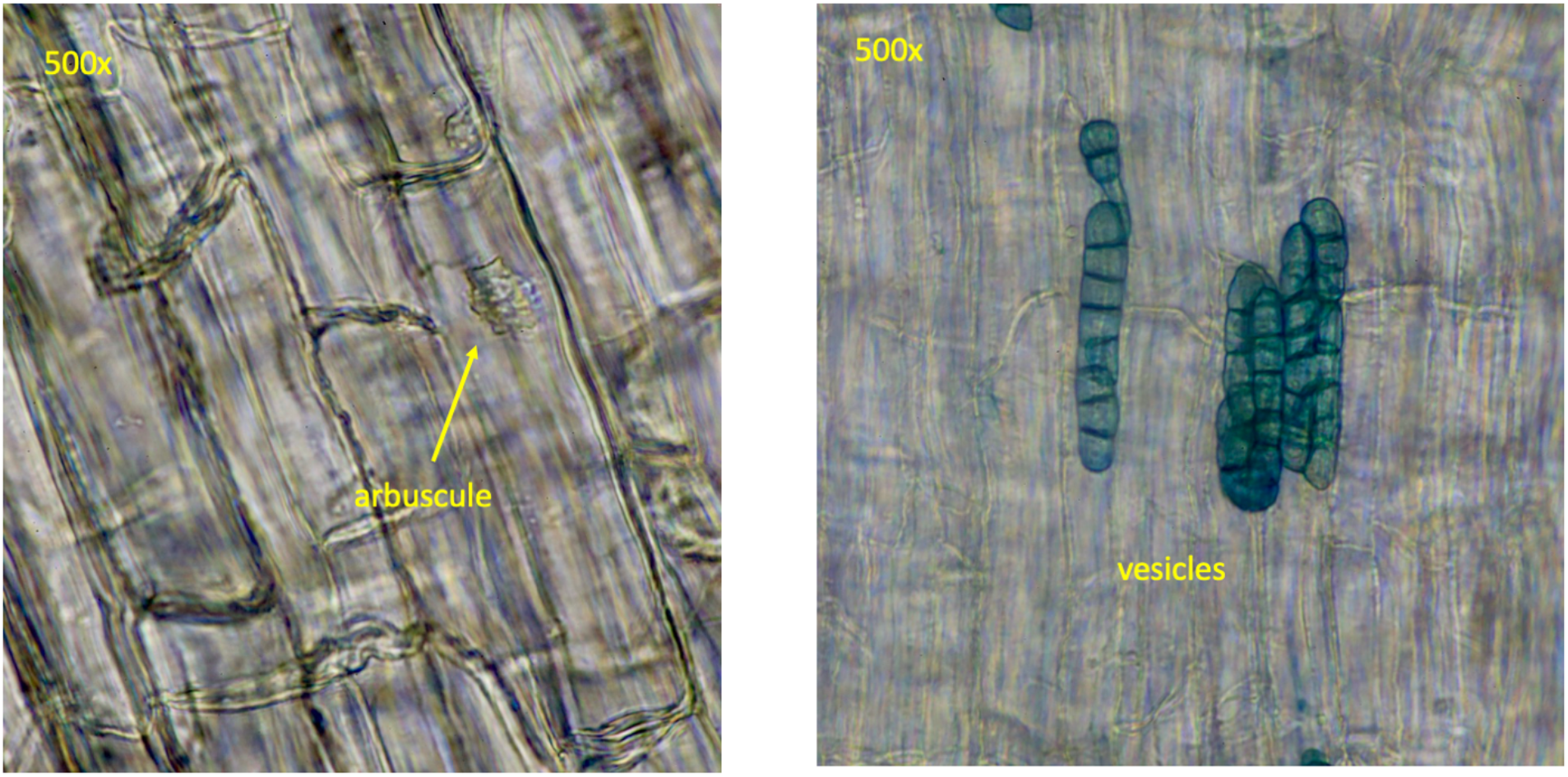
Compound microscope images (Olympus BX-60) of cleared and stained *Acer rubrum* root tissue showing the development of both arbuscules and vesicles typical of endomycorrhizal colonization (AM).

Dual colonization, or colonization of typically EM plants by AM type mycorrhizae is well documented in the literature. The potential for dual mycorrhizal colonization in tree seedlings has been noted for several tree species (Beauchamp, Stromberg, & Stutz, 2005; Chen, Brundrett, & Dell, 2000; Holste, Kobe, & Gehring, 2017; Moyersoen & Fitter, 1999), including several genera of *Quercus* (Egerton-Warburton & Allen, 2001; Holste et al., 2017; Querejeta, Egerton-Warburton, & Allen, 2009; Rothwell, Hacskaylo, & Fisher, 1983; Watson, Von Der Heide-Spravka, & Howe, 1990) as well as specifically in *Quercus rubra* (Dickie, Koide, & Fayish, 2001; Dickie, Koide, & Steiner, 2002b; Grand, 1969; Henry, 1934).

In a similar greenhouse study of tropical tree seedlings (Holste & Kobe, 2017) also found that most differences in growth were explained almost exclusively by plant species rather than mycorrhizae. Subjects included *Quercus costaricensis* and *Eucalyptus grandis*, known for associating with both EM and AM. In their experiment, four nutrient treatments were administered, inoculant was produced from field soil, and plants were grown from seed for 2 months and then treated for 3 months. Similarly designed experiments to ours in the future would do well to germinate seedlings to better control early environmental variables. Colonization levels were lower in both species (~12%) than our study (>50% average). Similar levels of AM colonization to our study have been quantified in field samples of *Quercus rubra* (Dobson et al., 2020) with a mean value of 58% (correspondence with author). While it is assumed that *Quercus* species reliably associate with EM at maturity, more studies are needed to characterize this clearly documented tendency to form mutualisms with AM during development.

Distribution of AM forms varied in our two species (Table 3) with significantly more hyphae formation in *Acer rubrum* and higher vesicle formation in *Quercus rubra.* Notably, Low N:P treatments for both species under intraspecific competition had the two highest mean rates of colonization (Figure 6). As colonization by EM was lacking in our *Quercus* seedlings, we were not able to accurately test our original hypothesis-that EM species should comparatively thrive in low N availability and AM species should thrive in low P availability. Future studies would better control and quantify the type and number of mycorrhizal spores found in inoculants as well as more closely monitor changes from initial available N and P. As both species were colonized by forms of endomycorrhizas, it would be valuable to know whether AM colonization provides a different degree of benefit by species-i.e., lies in a different position on the mutualism/parasitism spectrum in *Acer* compared to *Quercus* seedlings.

**Figure 6.**
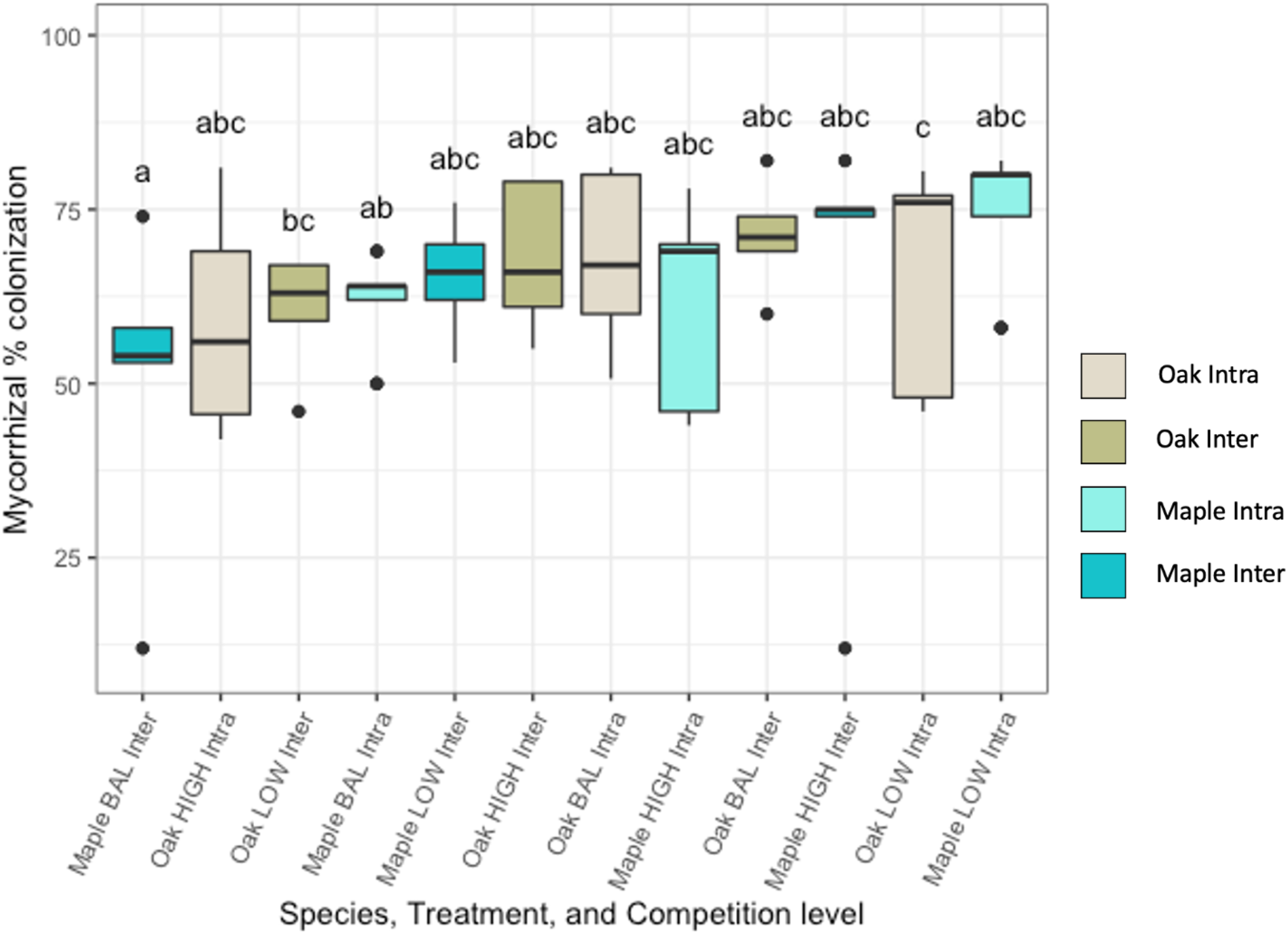
Boxplots of mycorrhizal colonization percentage by species and nutrient treatment when *Acer rubrum* and *Quercus rubra* seedlings are grown in both monoculture and in mixed competition for 3 months. Lower box boundary indicates the 25th percentile, the median value is shown as black line, and the upper boundary contains the 75th percentile. Error bars are whiskers above and below showing the 90th and 10th percentiles. Points outside the 90th and 10th percentiles (outliers) are also represented. Plots with the *same letters* above each box do not differ significantly by ANOVA (*p* < 0.05) among treatment groups.

Our results support the contention that intraspecific competition generally has a significantly stronger effect than interspecies competition on growth and development (Adler et al., 2018), though RGR varied more by competition level in heights and more by species in stem diameter measurements. Considering the number of studies that find evidence of colonization by AM in normally EM associating seedlings (Chen et al., 2000; Dickie et al., 2001; Dickie, Koide, & Steiner, 2002a; Egerton-Warburton & Allen, 2001; Holste et al., 2017; Moyersoen & Fitter, 1999) further study into this phenomenon is merited. (Egerton-Warburton & Allen, 2001) suggest that there may be a successional progression in some species where AM trade-offs are worthwhile for young seedlings and EM trade-offs in terms of soil nutrients become more beneficial as plants mature. Our results show that dual colonization, or colonization with a species of mycorrhiza other than what is usually found in mature trees, is more common than previously thought. Further testing and experimental designs may better quantify how important this plasticity is during early seedling survival and stand initiation. The influence of mycorrhizal colonization on seedling roots under changing nutrient availability has been shown to be significant for accessing recalcitrant soil nutrients, but how much it can overcome nutrient limitation compared to the physiological functional traits of species or the influence of interspecific competition patterns remains to be investigated.

## Acknowledgements

Thank you to all the members of the Brodersen and Bradford labs at the Forest School at the Yale School of the Environment for advice and assistance in completing the design, implementation, and analysis of this experiment-especially Craig Brodersen, Mark Bradford, and Aleca Borsuk. Much thanks to Simon Queensborough and Andrew Muehleisen for statistical mentoring. From the Ashton lab, Mark Ashton, David Woodbury, and Danica Doroski provided a year full of helpful guidance. De-wei Li from the Connecticut Agricultural Experiment Station helped identify fungal morphology and proofread the text. Vanessa Beauchamp from Towson University, and Colin Averill from ETH Zurich provided valuable edits to the article in draft. Funding was provided by the Wildlife and Wildlands Fund at the Yale School of Forestry and Environmental Studies.

